# Virus-inclusive single cell RNA sequencing reveals molecular signature predictive of progression to severe dengue infection

**DOI:** 10.1101/388181

**Authors:** Fabio Zanini, Makeda L. Robinson, Derek Croote, Malaya Kumar Sahoo, Ana Maria Sanz, Eliana Ortiz-Lasso, Ludwig Luis Albornoz, Fernando Rosso Suarez, Jose G. Montoya, Benjamin A. Pinsky, Stephen R. Quake, Shirit Einav

## Abstract

Dengue virus (DENV) infection can result in severe complications. Yet, the understanding of the molecular correlates of severity is limited, partly due to difficulties in defining the peripheral blood mononuclear cells (PBMCs) that are associated with DENV in vivo. Additionally, there are currently no biomarkers predictive of progression to severe dengue (SD). Bulk transcriptomics data are difficult to interpret because blood consists of multiple cell types that may react differently to infection. Here we applied virus-inclusive single cell RNA-seq approach (viscRNA-Seq) to profile transcriptomes of thousands of single PBMCs derived early in the course of disease from six dengue patients and four healthy controls, and to characterize distinct DENV-associated leukocytes. Multiple genes, particularly interferon response genes, were upregulated in a cell-specific manner prior to progression to SD. Expression of MX2 in naive B cells and CD163 in CD14^+^ CD16^+^ monocytes was predictive of SD. The majority of DENV-associated cells in the blood of two patients who progressed to SD were naive IgM B cells expressing the CD69 and CXCR4 receptors and antiviral genes, followed by monocytes. Bystander uninfected B cells also demonstrated immune activation, and plasmablasts from two patients exhibited antibody lineages with convergently hypermutated heavy chain sequences. Lastly, assembly of the DENV genome revealed diversity at unexpected genomic sites. This study presents a multi-faceted molecular elucidation of natural dengue infection in humans and proposes biomarkers for prediction of SD, with implications for profiling any tissue and viral infection, and for the development of a dengue prognostic assay.

**Significance:** A fraction of the 400 million people infected with dengue annually progresses to severe dengue (SD). Yet, there are currently no biomarkers to effectively predict disease progression. We profiled the landscape of host transcripts and viral RNA in thousands of single blood cells from dengue patients prior to progressing to SD. We discovered cell-type specific immune activation and candidate predictive biomarkers. We also revealed preferential virus association with specific cell populations, particularly naive B cells and monocytes. We then explored immune activation of bystander cells, clonality and somatic evolution of adaptive immune repertoires, and viral genomics. This multi-faceted approach could advance understanding of pathogenesis of any viral infection, map an atlas of infected cells and promote the development of prognostics.

## Introduction

Dengue virus (DENV) is a major threat to global health, estimated to infect 400 million people annually in over 100 countries [1]. The four serotypes of DENV are transmitted by a mosquito vector. A licensed vaccine has shown limited efficacy and increased hospitalizations in children [2,3], and there are currently no approved antivirals available for dengue treatment. The majority of symptomatic patients present with dengue fever experiencing flu-like symptoms. Five to twenty percent of these patients progress to severe dengue (SD), manifested by bleeding, plasma leakage, shock, organ failure, and sometimes death [4,5]. Early administration of supportive care reduces mortality in patients with SD [6], however, there are no accurate means to predict which patients will progress to SD. The currently utilized warning signs to identify dengue patients at risk of progressing to severe disease are based on clinical parameters that appear late in the disease course and are neither sensitive nor specific. This promotes ineffective patient triage and resource allocation and continued morbidity and mortality [7–9].

DENV strains are classified in four clades called serotypes 1 to 4. The greatest risk factor for SD is previous infection with a heterologous DENV serotype, which can cause antibody-dependent enhancement (ADE) of the secondary infection [10–13]. Aberrant activation of cross-reactive T-cells may also play a role [14]. Biomarkers for early detection of SD based on molecular features of the patient’s blood have been proposed. These efforts have focused on two experimental techniques: (i) flow cytometry of fixed blood cell populations [15], and (ii) gene expression in bulk RNA extracted from blood or peripheral mononuclear blood cells (PBMCs) [16–19]. Although useful, these studies suffer from several limitations. The majority of these studies identified genes whose altered expression is associated with but does not precede the onset of SD and therefore cannot be used as prognostic biomarkers. From a technical standpoint, flow cytometry has a high throughput but is constrained to a few protein markers that are selected a priori, making it excellent for separating known, discrete cell populations but less appropriate for screening the complex, dynamic landscape of cell types, subtypes, and states characteristic of immune responses. Transcriptomics performed on bulk cell populations can screen thousands of genes but its resolution is limited, because it cannot capture tissue heterogeneity. Averaging the signal over various cell populations is confounded by changes both in abundances of cell types and activation states. Coupling fluorescence activated cell sorting (FACS) with single cell transcriptomics can potentially combine the advantages of both approaches [20]. It has also been challenging to identify DENV-associated and DENV-infected immune cells in humans. Since the composition of the immune system is complex and dynamic, we hypothesized that an unbiased, whole-genome approach would be more suitable than techniques based on a small number of markers to pinpoint the exact subset of cells associated with DENV in vivo and identify biomarkers of severity in distinct cell populations.

To facilitate identification of host cells containing viral RNA and characterization of their transcriptional response, we recently developed a virus-inclusive single cell RNA-seq (viscRNA-seq) platform [21]. We previously used this platform to characterize transcriptome dynamics of infection with flaviviruses in cultured human hepatoma (Huh7) cells [21]. We simultaneously quantified host cell mRNA and virus RNA abundance at the single cell level, thereby identifying proviral and antiviral factors that correlated with intracellular viral abundance [21]. A similar approach was used to monitor host response to Zika virus infection in neuronal stem cells [22]. Here, we coupled FACS with viscRNA-Seq to identify virus-associated cells from human patients and studied the molecular signatures preceding the development of SD infection. The use of antibodies against surface proteins during FACS enabled enrichment for specific cell populations. Moreover, since viscRNA-Seq requires no genetic manipulation of the cells of interest, this approach enabled high-resolution screening of the whole human transcriptome for changes in gene expression at the single cell level.

## Results

### High-dimensional profiling of single cells from dengue virus infected patients

We combined FACS with viscRNA-Seq to profile the host and viral transcriptomes in peripheral mononuclear blood cells (PBMCs) collected early in the course of natural dengue infection in humans. Blood samples were derived from four healthy control subjects and six DENV infected patients: two who experienced an uncomplicated disease course and four who subsequently progressed to SD (Fig. 1A-B). All subjects were prospectively enrolled to a cohort that we established in Colombia (“Colombia cohort”) (Supplementary Tables 1 and 2). Disease severity was classified on-site using 2009 WHO criteria upon presentation and discharge [23]. Patients were enrolled within 2-5 days after symptoms onset based on clinical presentation compatible with dengue or dengue with warning signs and positive NS1 antigen and/or anti-DENV IgM antibody. Notably, patients who presented with SD at the earliest visit were excluded. Whole blood and serum samples were obtained upon presentation. qRT-PCR [24] and serological assays confirmed the diagnosis of DENV infection and excluded other arboviral infections (including Zika and chikungunya) [25]. IgG avidity testing distinguished primary from secondary dengue (Supplementary Table 1) [25]. PBMC samples were isolated, stored and shipped in liquid nitrogen.

**Figure 1:**
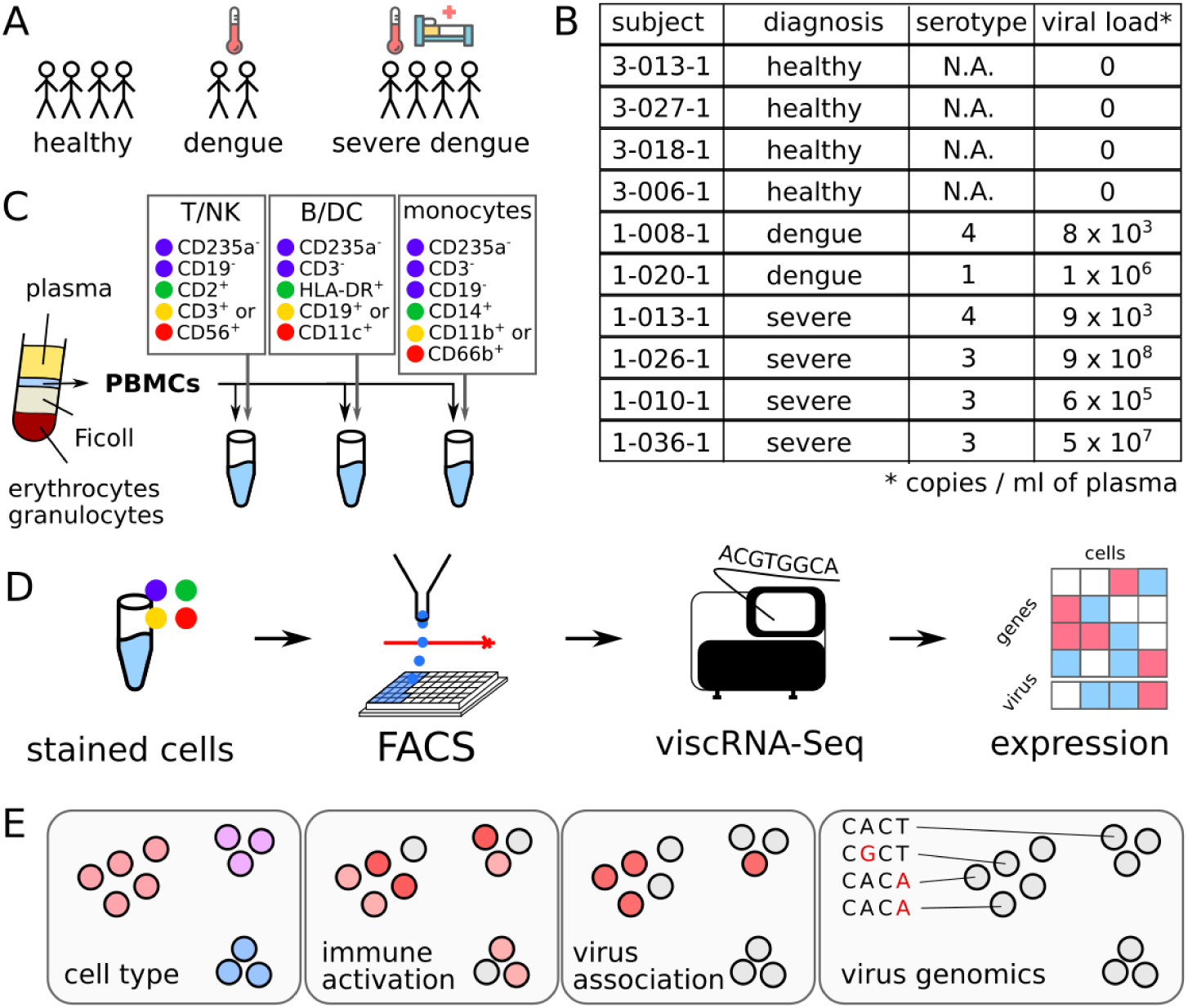
Fluorescence-assisted viscRNA-Seq workflow on peripheral mononuclear blood cells from DENV infected and healthy control human subjects. (A-B) Blood samples were collected from human subjects enrolled to the Colombia cohort (healthy, dengue, and severe dengue). (C) Peripheral blood mononuclear cells (PBMCs) were isolated via Ficoll centrifugation and stained with three antibody panels to distinguish various cell types: T, B, natural killer (NK), dendritic cells (DC), and monocytes. (D) Single cells from each aliquot were sorted and processed by viscRNA-Seq to simultaneously quantify single cell virus abundance and host transcriptome changes. (E) The information provided for each single cell includes: cell type, immune activation, infection state, and virus population genomics. N.A.= non-applicable.

To sort multiple types of immune cells in patient PBMC samples and enable viscRNA-Seq with high specificity and throughput, we assembled two panels of antibodies against host cell surface markers. The PBMC samples were split into several aliquots, immunostained (Fig. 1C and Supplementary Figures 1-3 and Supplementary Table 3), and sorted via FACS into T cells, natural killer (NK) cells, B cells, monocytes, and dendritic (DC) cells. The viscRNA-Seq protocol was then followed, and each cell was sequenced at a depth of ~1 million reads on NextSeq 500 and NovaSeq (illumina) instruments (Fig. 1D). To measure intracellular DENV RNA abundance, we conducted viscRNA-Seq using the previously reported pan-DENV capture oligo [21]. The information provided by this approach on each individual cell included the cell type, immune activation state, infection state (whether and how much DENV RNA the cell contains), and sequence of the virus strain (Fig. 1E).

### FACS-assisted viscRNA-Seq captures multiple cell types and activation states

Most human tissues including blood present a skewed composition of cell types. Unbiased cell capture, as routinely done in microfluidics protocols (e.g. [26]), produces detailed data on the most abundant cell populations, but fails to represent rare cell populations. To overcome this limitation, we combined FACS with a plate-based protocol to capture immune cells from samples containing less than 1,000,000 cells (because cells are sorted directly into single wells) with high sensitivity (as assessed by CD45 expression), and adequate representation of various cell populations (Fig. 2A) [27,28]. In total, we sequenced over 13,000 cells, of which several hundred showed robust signal for DENV RNA (Fig. 2A). Following quality filtering, tens to hundreds of cells were analyzed for most cell types of each sample, for a total of ~8,700 cells (Fig. 2B). Within each cell type, multiple distinct overlapping immune cell subtypes and cell states were well represented in the dataset (Fig. 2C and Supplementary Table 4). In particular, within B cells alone we profiled many naive, IgM/IGD double positive cells as well as isotype switched cells. Most B cells formed a continuum of differentiation, but we also identified two additional clusters: the first expressed markers of plasmablasts and plasma cells, whereas the second showed high expression of TYROBP, a transmembrane signalling protein that has been implicated in B cell proliferation [29].

**Figure 2:**
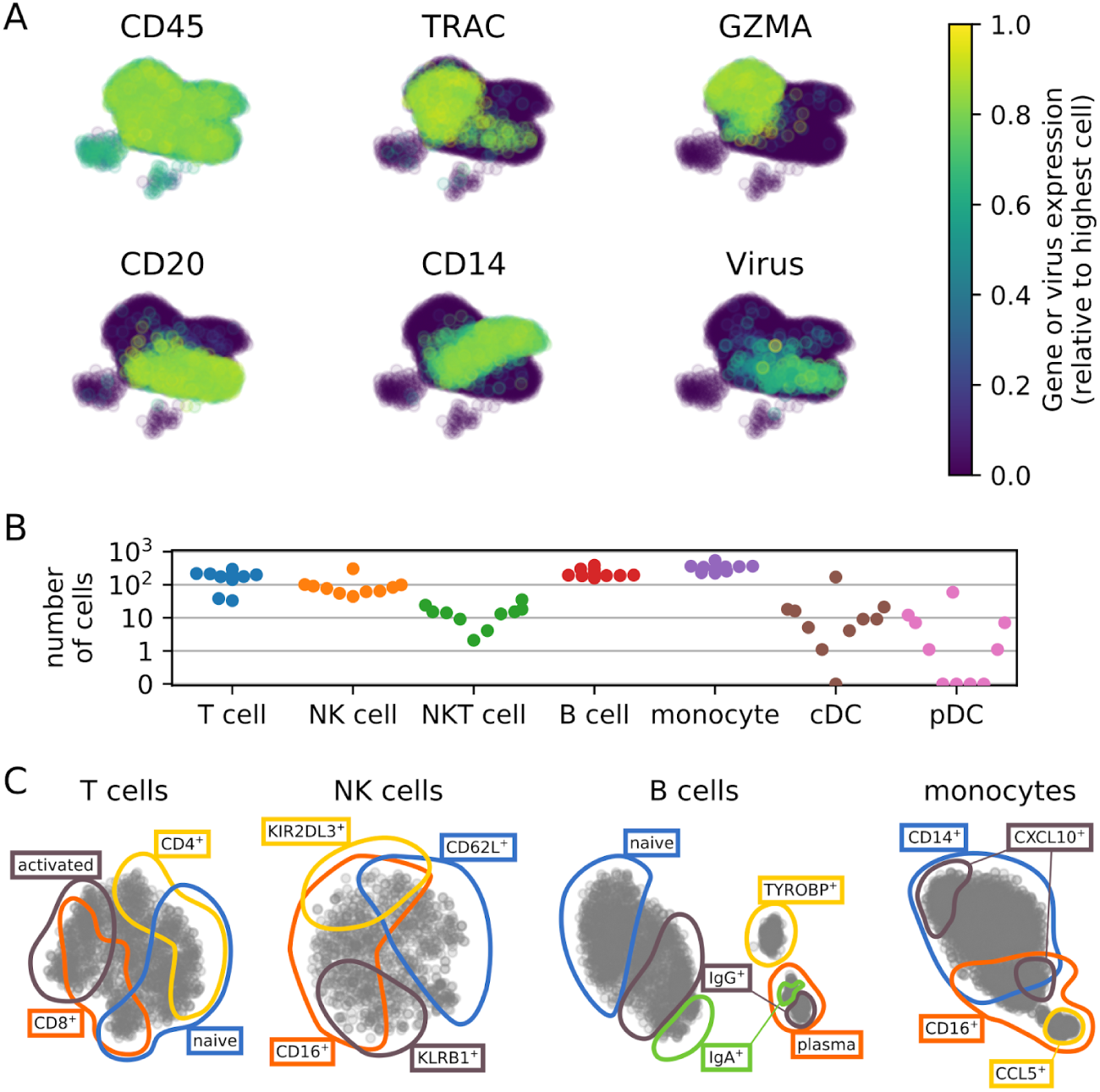
Overview of the types of peripheral mononuclear blood cells (PBMCs) surveyed. (A) Two-dimensional representation of the cells color coded by the expression level of cell type specific marker genes or the abundance of virus reads within the cell (> 30 virus reads per million reads in samples from two severe dengue patients, p1-026-1 and p1-036-1). (B) Number of cells analyzed for each cell type from each subject. (C) t-Distributed Stochastic Neighbor Embedding (tSNE) visualizations within T, NK, B cells, and monocytes, highlighting broad cell subtypes.

### Profiling single cell gene expression identifies candidate predictive biomarkers of severe dengue infection

Next, we profiled the host transcriptome responses in the various PBMC populations. Since blood samples were obtained early in the course of dengue infection, this analysis was aimed at revealing alterations in gene expression that preceded the progression to SD. For each cell subtype and gene, we compared the distribution of expression values across the three categories of subjects: healthy control (H); uncomplicated dengue (D), and severe dengue (SD). To identify differentially expressed genes, we used a two-sample Kolmogorov-Smirnov test together with a computation of fold change in the averages across cells. We identified several genes whose expression was strongly upregulated in subjects that subsequently progressed to SD. Many of these genes belonged to the antiviral interferon response, yet they were upregulated in a cell type specific manner (Fig. 3A). Some genes were expressed in multiple cell types but were upregulated more strongly in specific cells from SD subjects (Fig. 3B); other genes were expressed essentially only during SD except in a few cell types (Fig. 3C); a few genes were expressed only in one cell type and only in subjects who subsequently developed SD (e.g. CD163 in monocytes, Fig. 3D). These results indicate that distinct cell populations respond differently to the same viral infection, confounding the performance of bulk assays, such as microarrays. Since this heterogeneity is not a hindrance but rather a resource within the single cell approach, we then explored the predictive potential of gene expression in specific cell types. To do so, we averaged across cells within the same patient and cell population and tested binary classification of severity at increasing thresholds of expression, de facto simulating a pseudo-bulk assay that could be implemented in the clinic. We identified a number of genes in specific cell populations that showed great predictive power for distinguishing SD from other subjects, as assessed by receiver operating characteristic (ROC) curves (Fig. 3E). Two notable examples with optimal ROC performance (area under the curve = 1) are MX2 in naive B lymphocytes and CD163 in double positive CD14^+^/CD16^+^ monocytes.

**Figure 3:**
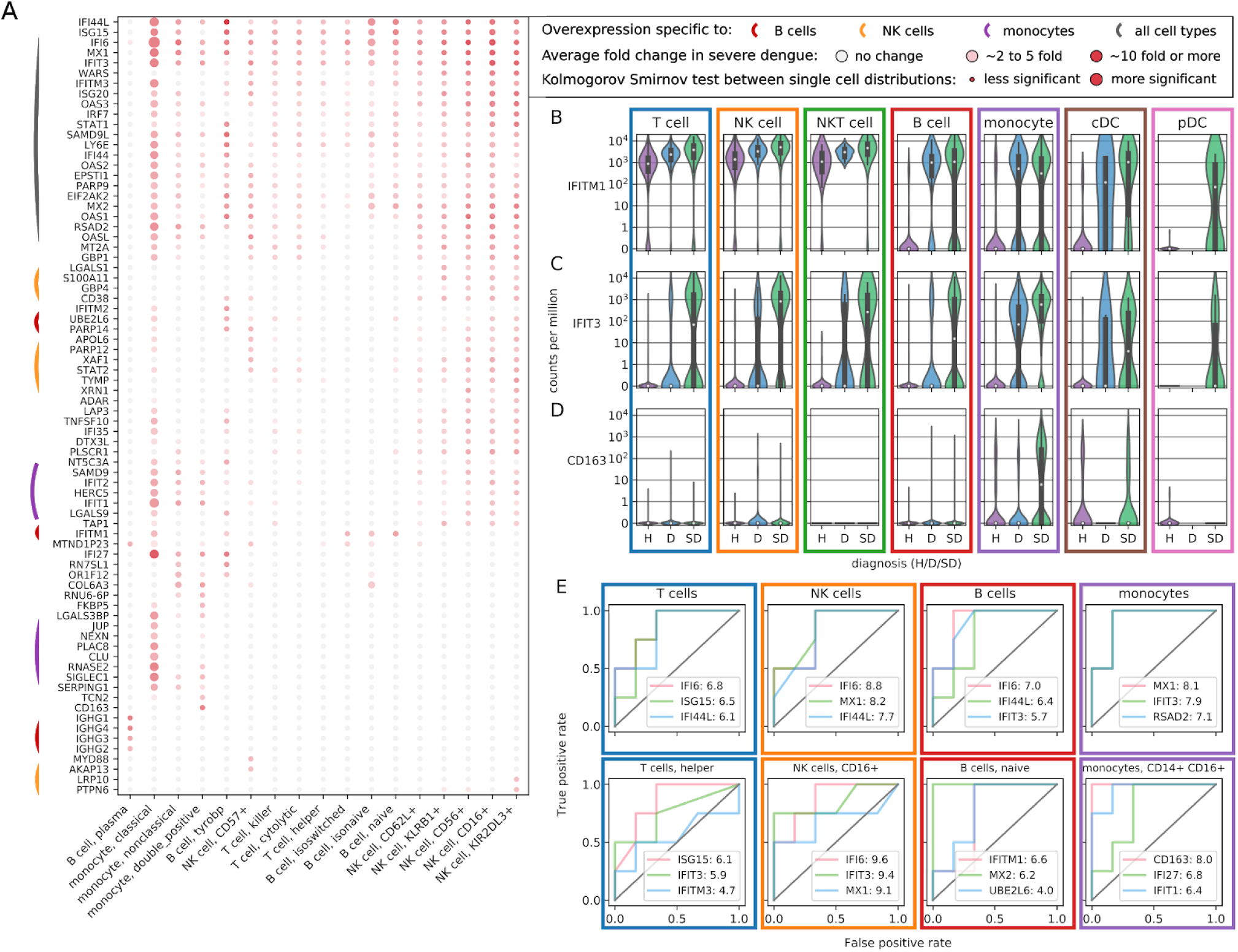
Differential expression across disease severity and cell types shows hallmarks predictive of severe dengue. (A) Genes that are overexpressed in subjects prior to progressing to severe dengue across cell types and subtypes. Color (white to red) indicates the average logfold change; size of the dot indicates lower P-value in a distribution statistical comparison (2 sample Kolmogorov-Smirnov). (B) Many inflammatory genes such as IFITM1 are expressed ubiquitously during both mild and severe dengue infection. (C) Other genes such as IFIT3 are specifically expressed prior to the development of severe dengue in various types of lymphocytes. (D) A number of genes show double specificity for both severe dengue and a single cell type, for instance CD163 in monocytes. (E) Averaging across cells within specific cell types and subtypes indicates promising candidate predictors of severe dengue as assessed by ROC curves at increasing discriminatory thresholds for gene expression versus disease severity. The numbers after the gene name indicate log2 fold changes of average expression in patients progressing to severe dengue versus other dengue patients, indicating an overexpression of these genes by a hundred fold or more in our cohort. H: healthy subject, D: dengue, SD: severe dengue.

### Virus in severe dengue patients is primarily associated with naive B cells

To define the cell subtypes that are associated with DENV in our PBMC samples, we then focused on cells with viral RNA reads. We detected viral reads in two samples only (out of six dengue confirmed samples analyzed), both of which were derived from subjects who had high viral loads in their serum and that subsequently progressed to SD (samples 1-026-1 and 1-036-1, Supplementary Table 2). In both samples, a small number of monocytes were associated with viral RNA. A weak upregulation of CD4 and other genes was observed in these virus-associated monocytes (Supplementary Fig. 4). The majority of virus-associated cells were B lymphocytes (Fig. 4A). No viral reads were detected in other types of leukocytes. These findings are in line with a previous report based on bulk qPCR assays [30]. The fraction of uniquely mapped reads corresponding to viral RNA in those cells was heterogeneous but generally 1% or less, corresponding to several hundred reads per cell but much lower than we measured in cultured Huh7 cells [21].

**Figure 4:**
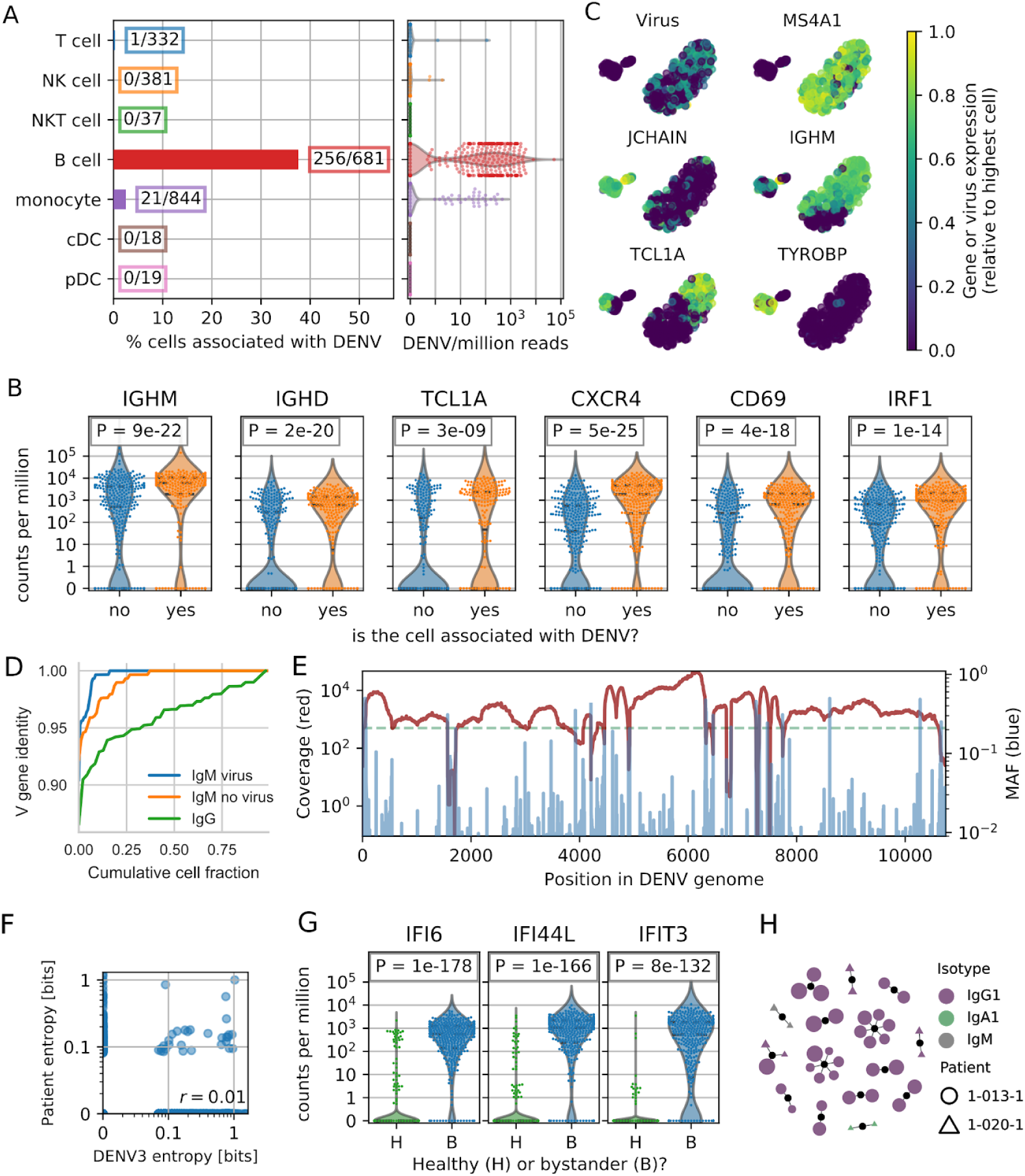
Dengue virus in two severe dengue patients is mostly associated with naive B cells. (A) Fraction of DENV-associated cells across cell types from the two subjects and relative amount of virus RNA from each cell. (B) Virus-associated B cells from the same subjects show a higher expression of specific surface receptors (CXCR4, CD69) and immune activation genes (IRF1, FCRL1). (C) tSNE visualization of the B cells from the two subjects. The expression level of DENV RNA and MS4A1 (CD20), JCHAIN, IGHM, TCL1A, and TYROBP are highlighted. (D) Fractional identity of heavy chain V loci to their germline counterparts in virus-associated IgM, bystander IgM, and IgG B cells from the subjects 1-026-1 and 1-036-1. (E) Coverage (red) and minor allele frequency (MAF, blue) along the DENV genome in the viral reads from all cells from patient sample 1-026-1 show the genetic diversity of the virus population. (F) Site specific Shannon entropy of a cross-sectional DENV serotype 3 alignment does not correlate with entropy from the viral reads of patient sample 1-026-1. Only sites with a coverage of 500 or more reads are considered (dashed green line in panel E). (G) B cells that are not associated with DENV (bystanders) but derived from subjects with virus-associated cells (B, blue) show a clear interferon response compared with B cells derived from healthy controls (H, green). (H) Graph of heavy chain CDR3 antibody clonality showing clonal expansion of IgG1 plasmablasts in patients 1-013-1 and 1-020-1. Each dot is a unique antibody sequence, larger size corresponds to more somatic hypermutation.

To determine whether a distinct subpopulation of B cells was specifically associated with DENV, we identified the most upregulated genes in the virus-associated population versus other B cells from the same patients. DENV-associated B cells were enriched but not exclusive to IgM/IGD isotypes as well as other markers of naive B lymphocytes, such as the transcription factor TCL1A (see Supplementary Fig. 5 for more markers) [31]. The surface receptors CD69, FCRL1, and CXCR4 that signal B cell activation and tissue-specific homing, and IRF1 that encodes an interferon related protein were also upregulated (Fig. 4B). We computed a 2-dimensional embedding of the B cells via t-Distributed Stochastic Neighbor Embedding (tSNE) from the two PBMC samples with detectable viral reads and measured no viral reads associated with cells belonging to the plasma cells or TYROBP^+^ clusters (Fig. 4C and Supplementary Fig. 5) [32]. We then assembled the whole B cell receptor (BCR) locus *de novo* and found that virus-associated, IgM B cells tended to have less hypermutations than other IgM B cells from the same subjects (Fig. 4D). In contrast, V/J usage in heavy and light chains was not apparently different between virus-associated and bystander B cells in the same subjects (Supplementary Fig. 6). Moreover, the expression of a number of genes that we have shown correlated with intracellular viral abundance in DENV-infected Huh7 cells and are known to participate in intracellular viral dynamics was not altered in the virus-associated B cells, raising the possibility that these cells are associated but not infected with DENV (Supplementary Fig. 7). We performed single molecule fluorescence in situ hybridization (smFISH) to detect positive and negative strand DENV RNA from naive B cells and monocytes from a healthy blood donor. We observed no evidence of either DENV strand in these B cells, unlike monocytes or control Huh7 cells (Supplementary Fig. 8).

In addition to counting the DENV reads, we mapped them in an iterative manner and recovered ~300,000 viral reads from patient 1-026-1 and ~2,000 reads from patient 1-036-1. We obtained high coverage across the whole DENV genome and a third of the genome, respectively. The intrapatient population genomics showed a wide range of conservation levels, as determined by minor allele frequencies (Fig. 4E and Supplementary Fig. 9A). Site-specific Shannon entropy restricted to positions with 200 or more virus reads did not correlate with cross-sectional entropy in DENV serotype 3 (Fig. 4F and Supplementary Fig. 9B).

Hundreds of non DENV-associated B cells (henceforth called “bystanders”) were recovered from samples containing DENV-associated cells. We computed differential gene expression between these bystanders and B cells from healthy controls and identified a strong antiviral response via interferon stimulated genes IFI6, IFI44L, and IFIT3 (Fig. 4G). Moreover, we considered whether the diversity of the immune repertoire (B and T cell receptors - BCR and TCR) could play a role in virus-cell association. Whereas assembled BCRs from patients with detected DENV-associated B cells scattered into small clones, the BCR repertoire of patients 1-013-1 and 1-020-1, who had no DENV-associated B cells, contained large clonal families comprised of multiple plasmablasts sharing similar antibody heavy chains, indicating a rapid and large clonal expansion in the B cell compartment (Fig. 4H). The fact that such plasmablast expansions were captured simply as part of these patients’ circulating B cell populations was surprising given the vast diversity of possible BCR rearrangements [33] and could be indicative of a more extensive plasmablast response and concurrent rise in neutralizing antibody titers known to occur in response to acute dengue infection [34]. One clonal family had members belonging to both patients, while another featured two plasmablasts with nearly identical heavy chains, but distinct light chains, raising the possibility of parallel somatic evolution [35]. Within the T cell compartment, we found that clustering by TCRβ/δ CDR3s produced clonal families that were largely private to an individual, while clustering according to TCRα/γ CDR3s revealed known invariant T cell subsets, including invariant natural killer T cells (iNKT) and mucosal associated invariant T cells (MAIT), as well as public γ chain CDR3 sequences (Supplementary Fig. 10) [36].

## Discussion

We recently developed viscRNA-Seq, a scalable approach to quantify host and non-polyadenylated viral RNAs from the same cell [21]. In the current study, we apply viscRNA-Seq to in vivo samples for the first time and show that it can be used to effectively profile the landscape of host transcripts and viral RNA in thousands of single immune cells during natural dengue infection of human subjects. The human samples studied here posed additional challenges, beyond those presented in cultured cells. First, the exact sequence of the viral strain infecting the patient was unknown. We therefore designed oligonucleotide for virus capture in a conserved region of the viral genome [21]. Second, the target cell types of DENV in vivo are incompletely characterized, mandating assembly of antibody panels for FACS that maximize the probability of capturing the virus-associated cell population(s) (Fig. 1C). Third, since cell viability and integrity of the RNA after freezing, shipping, and thawing the PBMC samples was much lower than in cultured cells, we increased the throughput and PCR preamplification to ensure a sufficient number of high-quality cells from each cell type. These modifications have successfully addressed these challenges, as indicated by our findings.

Because the viscRNA-seq approach can be extended to any virus of interest and is compatible with surface markers for FACS, we expect it to be readily extendible to dissociated solid tissues, for instance to characterize viral reservoirs of various viral infections. A similar approach was recently applied to influenza infection in mouse lungs [37] and to in vitro Zika virus infection of neuronal stem cells [22].

There are currently no clinically usable biomarkers to predict the development of severe complications associated with DENV infection including bleeding, shock, vascular leakage, and organ failure [4]. Previous work on molecular biomarkers for dengue severity has focused on flow cytometry, which has high throughput but requires an a priori choice of a few tens of marker proteins [15], and bulk transcriptomics, which can quantify the expression of all genes in parallel but is confounded by the superimposition of cell types (e.g. B and T lymphocytes) and cell state (activation of SD specific genes) [16,17]. Since obtained prior to the progression to SD, the blood samples analyzed in this study, combined with the single cell resolution and ability to sample a wide range of cell types and activation states via viscRNA-seq, provided a unique opportunity to discover predictive biomarkers of severity. Our data suggests the expression of MX2 within naive B cells and of CD163 within CD14^+^ CD16^+^ monocytes is greaty upregulated prior to the development of SD (Fig. 3E). It has been previously reported that MX2 is one of only four interferon-induced genes induced in an IRF3 and IRF7 independent manner in DENV infected mice [38], and that CD163 in macrophages and CD14^+^ CD16^+^ monocytes contributes to the pathogenesis of SD [39,40]. Given the small number of subjects analyzed in this study, the predictive power of these candidate biomarkers warrants further validation in larger cohorts. Nevertheless, these findings underscore the utility of the viscRNA-Seq approach to identify prognostic biomarkers for dengue.

Cell lines such as Huh7 and primary cells such as monocyte-derived dendritic cells are commonly used to study DENV-host cell interactions, however, elucidating the association of DENV with human cells in vivo has proven challenging. To the best of our knowledge, this is the first study to perform high-dimensional profiling of DENV-associated cells in vivo. In agreement with the expectation that myeloid cells are more easily infected than other cell types [11], we found 21 monocytes containing viral RNA in two patients with high viral load prior to the progression to SD. We observed a weak upregulation of CD4 and other genes in these virus-associated monocytes (Supplementary Fig. 4). Many more B cells from the same patients contained viral RNA, in line with a previous report on bulk samples [30]. Distribution-level statistics and dimensionality reduction indicated that IgM positive, naive B cells expressing the surface markers CD69 and CXCR4 are the subset of PBMCs that most commonly associate with DENV in vivo, although other B cells of other isotypes were also represented. In contrast, plasmablasts, plasma cells, and TYROBP positive B cells had no viral reads (Fig. 4A-D). B cells are known to be involved in SD pathogenesis by producing antibodies for antibody dependent enhancement (ADE) [12–14]. Discovering that naive B cells possessing diverse B cell receptor (BCR) sequences, rather than specific isotype switched, sequence-restricted memory B cells, are associated with DENV was therefore surprising. More work is needed to conclusively define whether these B cells are infected with DENV or just associated with it. Nevertheless, our data thus far based on lack of alteration in expression of genes known to be altered in DENV infection and the absence of DENV positive B cells in PBMCs via FISH support the latter. It is possible that these virus-associated B cells shuttle DENV within the human body. Bystander B cells from patients with detected DENV-associated cells had elevated levels of immune genes, particularly of the interferon response (Fig. 4G). B cells from one severe dengue and one dengue patient samples (1-013-1 and 1-020-1) showed an interesting clonal structure in terms of antibody sequences. Specifically, multiple antibody sequences of heavy and light chains from several cells from patient 1-013 clustered into a few, presumably very large lineages of mostly heavily hypermutated IgG1 plasmablasts. Since no DENV RNA reads were detected in these samples (in contrast to samples 1-026-1 and 1-036-1), we speculate that this oligoclonal plasmablast population may be reducing binding of DENV by the host B cells. Further work is needed to understand the role of virus-associated and bystander B cells, determine whether DENV prevents the hypermutation upon binding naive B cells, and decipher whether the detected large antibody lineages are protective against DENV itself.

From the viral reads of two patients, 1-026-1 and 1-036-1, we assembled the entire or a third of the DENV genome, respectively, and observed some high-variability genomic sites (Fig. 4E). Previous work on other RNA viruses, particularly HIV-1, has shown that due to error-prone viral polymerases and fast generation times, intrapatient genomic viral diversity can represent a subsampled snapshot of the global diversity of the same virus in multiple infected individuals, implying a universal landscape of fitness costs [41,42]. DENV behaves quite differently, as globally variable sites do not correspond to variable sites within our patients (Fig. 4F). An optimized approach with higher sensitivity and sample selection (PBMCs or solid tissues) that maximizes the number of viral reads will facilitate a deeper understanding of the genomic diversity of viruses inhabiting the human body at the single cell level.

In this study, we leveraged the viscRNA-Seq approach to explore many different facets of virus infection in uncomplicated and severe dengue in humans at the single cell level. This multi-faceted profiling included investigation of transcriptional upregulation in specific subpopulations as a predictor of disease severity. Further validation in larger cohorts is warranted to determine the effectiveness of the identified candidate biomarkers as potential prognostic tools. Cell purification (e.g. by magnetic beads) followed by a rapid bulk expression assay (e.g. qPCR) is one option to translate such findings into a near-care, sample-to-answer system assay to be used for predicting progression of SD upon patient presentation. We also explored preferential association of virus with certain host cells, immune activation of bystander cells, clonality and somatic evolution of the adaptive immune repertoire, and intrapatient viral genomics. This technological convergence, combined with a high level of experimental and computational automation, underscores the utility of viscRNA-Seq as a powerful tool to rapidly gain a broad knowledge of emerging infectious diseases from just a few tissue samples.

## Acknowledgements

This work was supported by seed grants from the Stanford Bio-X Interdisciplinary Initiatives Seed Grants Program, the Stanford Translational Research and Applied Medicine (TRAM) program, the Stanford SPARK program, Stanford Child Health Research Institute, and Stanford Institute for Immunity, Transplantation, and Infection to S.E. This work was also supported by NIH (5U19AI057229-15), by the Chan Zuckerberg Biohub, and by the California Institute for Regenerative Medicine (GC1R-06673) to S.R.Q. F.Z. was supported by a long-term European Molecular Biology Organization (EMBO) fellowship (ALTF 269–2016). M.R. was supported by the Stanford Advanced Residency Training at Stanford (ARTS) Fellowship Program. We are thankful to the patients who participated in this study and to their families.

